# Normalization of RNA-Seq Data using Adaptive Trimmed Mean with Multi-reference

**DOI:** 10.1101/2023.12.04.570016

**Authors:** Vikas Singh, Nikhil Kirtipal, Byong-Sop Song, Sunjae Lee

## Abstract

The normalization of RNA sequencing data is a primary step for downstream analysis. The most popular method used for the normalization is the trimmed mean of M values (TMM) and DESeq. The TMM tries to trim away extreme log fold changes of the data to normalize the raw read counts based on the remaining non-deferentially expressed genes. However, the major problem with the TMM is that the values of trimming factor M are heuristic. This paper tries to estimate the adaptive value of M in TMM based on Jaeckel’s Estimator, and each sample acts as a reference to find the scale factor of each sample. The presented approach is validated on SEQC, MAQC2, MAQC3, PICKRELL, and two simulated datasets with two groups and three groups conditions by varying the percentage of differential expression and the number of replicates. The performance of the present approach is compared, and it shows better in terms of area under the receiver operating characteristic curve (AUC) and differential expression. The implementation of the present approach is available on the GitHub platform: https://github.com/vikkyak/Normalization-of-Bulk-RNA-seq.

## Introduction

High throughput RNA-seq is one of the most effective tools for investigating various biological and medical applications. Highly complex and massive data sets generated by sequencers initiate a need to develop statistical and computational data analysis methods (Zyprych-Walczak et al. [2015]). In RNA-Seq, the RNA is fragmented and reverse-transcribed to cDNA or vice-versa. These fragments are sequenced, and produce reads aligned to a pre-sequenced reference genome or transcriptome or, in some cases, assembled without the reference. These reads are mapped to a gene and used to quantify its expression (Evans et al. [2018]). The raw read counts have a different source of systematic variation, which includes differences between-sample such as library size (Mortazavi et al. [2008]) or differences within-sample, gene length (Oshlack and Wakefield [2009]), and guanine-cytosine (GC) content (Risso et al. [2011]). These variations affect the differential expression (DE) analysis of the RNA-seq data. It can be overcome by a suitable normalization method similar to microarray-based gene expression data analysis (Singh et al. [2019], Singh and Verma [2022]). The authors (Park et al. [2003], Bullard et al. [2010]) have described how normalization affects the differential gene expression analyses in microarray data. Arguably, the choice of normalization method can significantly affect the downstream analysis results more than the method used for completing differential expression (Park et al. [2003]).

The normalization is classified into two categories, i.e., within-sample and between-sample. Within-sample normalization helps to correct the expression level of each gene associated with the expression level of other genes in the same sample. The most broadly studied methods for within-sample and between-sample normalization are Reads Per Kilobase Million (RPKM) (Mortazavi et al. [2008]) and Fragments Per Kilobase Million (FPKM) (Trapnell et al. [2010]). However, the different gene lengths can lead to a bias in per-gene variance for differential expression analysis of low abundance genes (Mortazavi et al. [2008]). In the literature, within-sample normalization approaches for RNA-seq data have corrected the biases arising from library size, gene length, and GC content. These normalization methods are Total Counts (TC) (Risso [2011]), Upper Quartile (UQ) (Bullard et al. [2010], Robinson et al. [2010]), per-sample Median (Med) (Bullard et al. [2010]), DESeq normalization (Love et al. [2014]), Trimmed Mean of M values (TMM) (Robinson and Oshlack [2010]), Tag Count Comparison (Kadota et al. [2012]) and sequencing data based on a Poisson log-linear model (Li et al. [2012]). To correct the library size, TC, UQ, Med, DESeq, and TMM are used, and they are based on a common normalizing factor per sample to normalize the genes. Selecting the optimal method from the perspective of sensitivity and specificity will be challenging due to biological variation, read depth and the number of biological replicates in the RNA-seq data (Sun and Zhu [2012]). Based on the DE analysis, the previous studies suggested that the DESeq and TMM perform better than the rest of the methods (Kvam et al. [2012], Soneson and Delorenzi [2013], Seyednasrollah et al. [2015]).

In bulk RNA-seq, the data may vary based on various uncontrollable experimental conditions. RNA-seq raw data must be normalized to scale the read counts. The most widely used and well-accepted methods for bulk RNA-seq data analysis are the TMM and DESeq (Li et al. [2017]). In the TMM, trimmed factor values are heuristic or user-defined. In this paper, we have presented an adaptive approach that selects the trimming factor from data automatically. The value of the trimming factor is estimated from the data using Jaeckel’s estimator, which helps to find a more robust factor by minimizing the asymptotic variance estimate of the alpha-trimmed mean. Additionally, in the TMM, only one sample from the data acts as a reference signal to find the value of the scale factor, which may be biased. To overcome this effect, we have used all the samples as reference signals to calculate the scale factor.

The rest of the paper is organized into four sections. Section 2 discusses the present method. Experimentation and performance evaluation are described in Section 3. while Section 4 briefly concludes the paper.

The key contributions of the paper are briefly described as:

- The value of the trimming factor is estimated from the data using Jaeckel’s estimator, which offers a robust trimming factor to trim away the extreme log fold changes. The trimming factor is obtained by minimizing the asymptotic variance estimate of the alpha-trimmed mean estimator.
- TMM uses only one sample from the data as a reference signal to find the scale factor, which may be biased. To overcome this effect, we have used all the samples as reference signals to determine the scale factor. We get the scale factor matrix of order *n* * *n* and apply the geometric mean corresponding to the column to find the common scale factor.

## Proposed Approach: Adaptive Trimmed Mean of M

As described (Robinson and Oshlack [2010]), the trimmed mean is the average after trimming the upper and lower values (x%) of extreme log fold changes. The TMM method is dual-trimmed by log fold changes (*M*_*g*_) (*k*^*th*^ sample relative to *r*^*th*^ sample for gene g) and by absolute expression intensity (*A*_*g*_). The default trimming factor for *M*_*g*_ is 30 %, and for *A*_*g*_ is 5% (Robinson and Oshlack [2010]). The gene-wise log fold changes and absolute expression intensities for sequencing data are defined as follows:

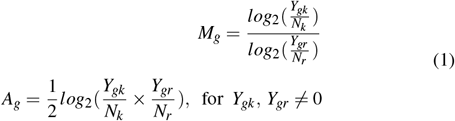

where, *Y*_*gk*_ and *Y*_*gr*_ be the observed read count of gene *g* with respect to sample *k* and reference sample *r, N*_*k*_ and *N*_*r*_ be total number of raw read counts for sample *k* and *r*, respectively.

To robustly summarize the observed *M*_*g*_ values, the authors have trimmed both the *M*_*g*_ and *A*_*g*_ values before taking the weighted average. The weights are utilized to account for the fact that log fold changes from genes with large read counts have small variance on the logarithm scale (Robinson and Oshlack [2010]). As explained, the normalization factor for sample *k* using reference sample *r* is determined as:

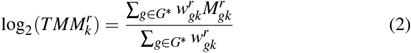

Where

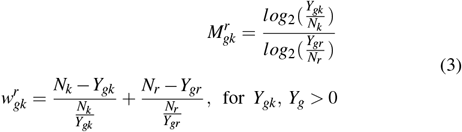

where 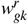 is the weight as the inverse of the asymptotic variance.

The TMM performs well, but the key issue is selecting the optimal trimming factor value. To get the optimal values of trimming factor we have used the asymptotic properties of Trimmed Means.

Let *X*_1_, *X*_2_, …, *X*_*n*_ be a sample of independent, identically distributed (iid) random variables with a common symmetric distribution. The *X* maybe the *M*_*g*_ or *A*_*g*_ values. The alpha-trimmed mean is given as

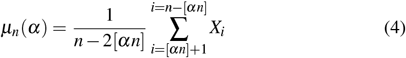

With the assumption that *E*^−1^(*α*) and *E*^−1^(1 − *α*) are unique, it is shown (Stigler [1973], Oten and de Figueiredo [2004]) that Equation (4) is an asymptotically normal estimator with respect to sample asymptotic variance estimate (*V* (*α*)), i.e.,

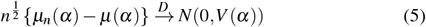

where

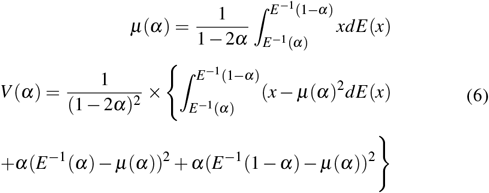

The asymptotic alpha-trimmed mean estimator (*μ*(*α*)) in Equation (4) is optimized by selecting an *α*_*opt*_ such that,

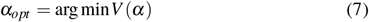

The continuous from of asymptotic variance can be written in discrete form using the Jaeckel’s Estimator as follows:

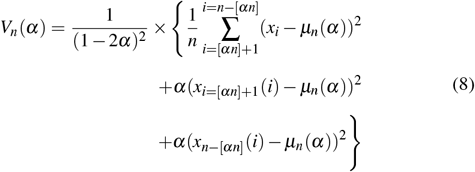

The optimal values of trimming factor (*α*) is obtained as

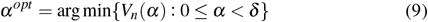

The default value of *δ* is 0.5 and the proposed approach is graphically described in Fig. 1.

**Fig. 1.**
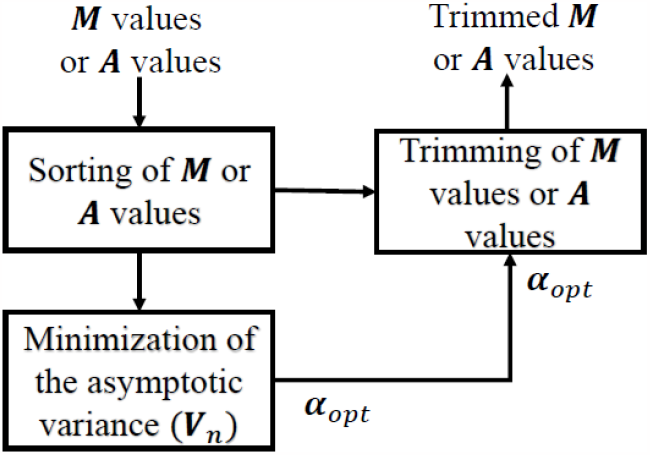
Pictorial representation of learning *α*_*opt*_ using Jaeckel’s estimator for trimming of *M* or *A*.

### Algorithm 1 Robust Estimation of Trimming Factor (*α*)

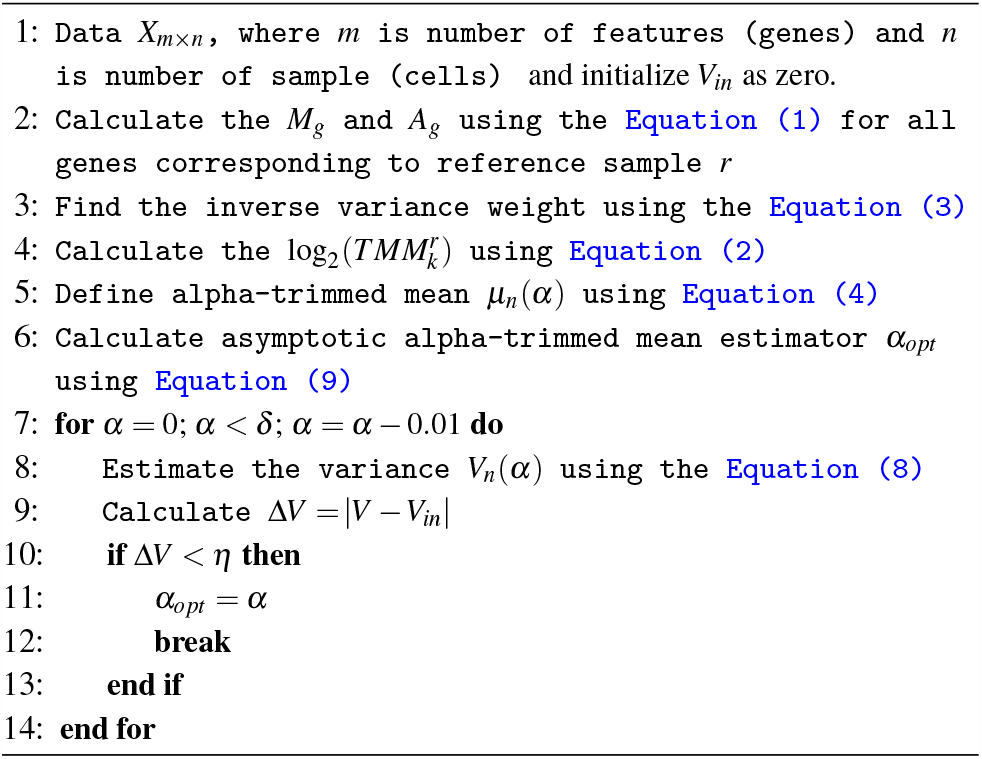

## Results & Discussion

This section described the detailed description of datasets and performance evaluation of the normalization methods.

### Real Datasets

#### Sequencing Quality Control (SEQC) Dataset

The performance of normalization methods is examined on the RNA-Seq dataset of the SEQC project (Su and Mason [2014]). This project described RNA-Seq technology across distinct platforms and alignment methods and collected the dataset of four samples, i.e., A, B, C, and D, with the number of replicates per sample (see Su and Mason [2014]). The SEQC dataset also has TaqMan qRT-PCR measurements on 1000 genes. The PCR data are commonly used to identify true differential gene expression and determine false negatives and positives in RNA-Seq data. The complete SEQC qRT-PCR data have 1044 genes. Here, we also perform a similar study as presented (Evans et al. [2018]), in which the PCR data is matched with SEQC RNA-Seq data, selected common genes with enough information, and eliminated duplicate genes. The unique genes obtained from both RNA-Seq and PCR measurements are 733 genes.

#### Microarray Quality Control Project (MAQC) Dataset

In the second study, we analyzed the performance of each method on the MAQC2 and MAQC3 datasets of the MAQC Project (Shi et al. [2006]). The MAQC2 has two RNA-Seq datasets from the MAQC project with two distinct biological samples, i.e., human brain reference RNA (hbr) and universal human reference RNA (uhr). The MAQC2 is accessed from the NCBI sequence read archive with reference ID SRX016359 (hbr) and SRX016367 (uhr), and it consists of a read length of 36bp (Bullard et al. [2010]) and second dataset (GEO series GSE24284) consists of the 50bp hbr (sample ID: GSM597210) and uhr (sample ID: GSM597211) RNA samples (Wan and Sun [2012]). MAQC3 is accessed from GEO (GSE49712), and it has five technical replicates in two biological conditions (uhr and hbr) (Rapaport et al. [2013]).

#### Pickrell Dataset

Pickrell’s real data is accessed from the recount2 database with the sample ID “SRP001540”. It has an order of count matrix 58, 037 × 160 of human data generated from two sequencing centers, i.e., Yale and Argonne (Collado-Torres et al. [2017], Pickrell et al. [2010]). The dataset obtained from both center show similar results (Hänzelmann et al. [2013]), so we perform normalization on the Yale dataset with 79 samples. The column matrix of the dataset is reduced by summing the samples with technical replicates, which results in 69 samples and genes with a zero count value in all samples being removed, and the analyzed genes are 51,910. The analyzed Pickrell dataset consists of a count matrix 51, 910 × 69 that compares the expression levels of lymphoblastoid cells between 29 males and 40 females.

### Simulated Datasets

The effectiveness of the normalization methods is also validated on the simulated data by varying the proportion of DEGs and DEGs up-regulated in the individual conditions. The simulated data of two and three-group conditions are generated with the help of the simulateReadCounts function of TCC package in R (Sun et al. [2013]).

#### Two-group Comparison

The two groups simulated data consist of g = 10,000 genes. The number of simulated biological replicates of individual groups are *r*_1_ = *r*_2_ = 3, proportion of DEGs (PDEG = 0.25), DEGs up-regulated in the individual conditions are *P*_1_ = 0.9 (or *P*_2_ = 0.1), and degree of DEGs is fixed at four-fold (FC = 4).

#### Three-group Comparison

As described in Tang et al. [2015], we also validated on three-group simulated data with numbers of simulated biological replicates are *r*_1_ = *r*_2_ = *r*_3_ = 3. The simulated conditions are g = 10,000, the proportion of DEGs (PDEG = 0.25), and the degree of DEGs is fixed four-fold (FC = 4). Here, we generated two conditions for each group with the proportion of upregulated DEGs (1/3, 1/3, 1/3), and (0.4, 0.2, 0.4) (Osabe et al. [2021]).

### Performance Evaluation

#### Performance Analysis on Real Datasets

In this study, the proposed approach is compared with four widely used normalization methods, i.e., DESeq (Love et al. [2014]), PoissonSeq (Li et al. [2012]), DEGES (Kadota et al. [2012]) and TMM (Robinson and Oshlack [2010]) on real datasets. Here, we briefly describe the libraries used for the normalization and statistical tests. edgeR (Robinson et al. [2010]) performs TMM normalization based on the Bayes estimation and the exact test with a negative binomial distribution for DE genes analysis. DESeq2 uses median-based normalization to account for the presence of different library sizes. DESeq2 estimates the gene-wise dispersions and shrinks these estimates to generate more accurate dispersion estimates to model the counts. Finally, it fits the negative binomial distribution model and performs hypothesis tests using the Wald or Likelihood Ratio tests. TCC (Sun et al. [2013]) is a multi-step normalization method (called DEGES) that uses inbuilt functions of other libraries, i.e., DESeq2, edgeR, and so on. PoissonSeq, another software used in our analysis, is based on an iterative process. It estimates a group of non-DE genes, and the scaling factor of each sample is determined using the respective group. In this study, we have performed a Wald test to find the differentially expressed genes for DESeq2, DEGES, and PoissonSeq.

Genes with small read counts across the libraries have very little information for differential expression. These genes are filtered automatically by using the edgeR function filterByExpr. After removing the low-expressed genes, the data is normalized using different normalization methods. For the SEQC data, we have performed two tests, first for symmetric expression and the other for asymmetric expression, and performance is compared in terms of AUC values. As shown (Fig. 2), if the data using a complete set of 733 genes is normalized using the proposed approach, it performs better than the rest of the methods. However, in the case of asymmetric expression, the proposed method also performs a little better except for DEGES (Fig. 3). To analyze the performance of normalization methods under asymmetric expression, we have taken a subset of 619 PCR-validated genes with 75% of DE genes up-regulated in sample A, and the rest 25% of DE genes are up-regulated in sample B. We have also shown the false discovery rate (FDR) values (Table 1) in the case of two and five technical replicates. The area under the receiver operating characteristic curve (AUC) is used to compare the results of all methods based on the log-fold-change *<*= 0.5.

**Table 1.**
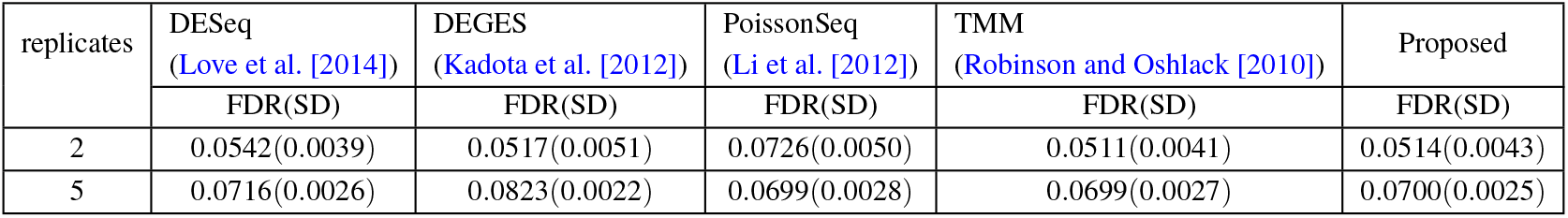
FDR with Standard Deviation (SD) for RNA-seq dataset of SEQC project with two and five replicates per Sample.

**Fig. 2.**
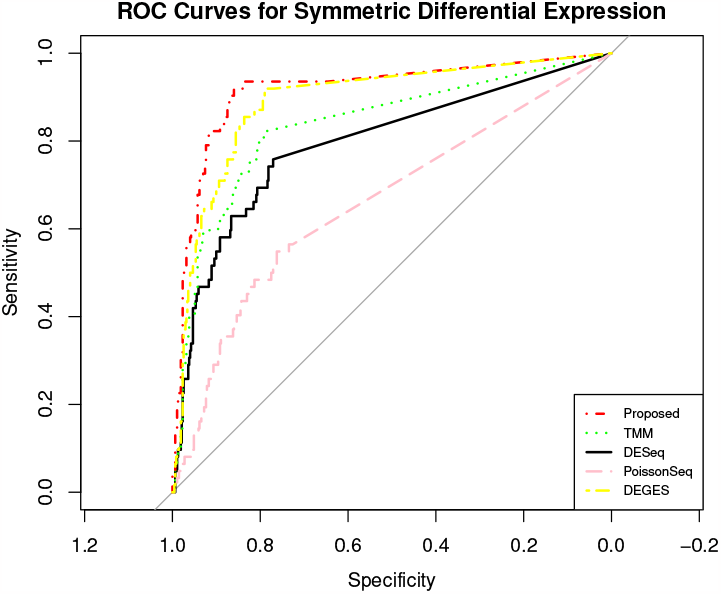
ROC curves for symmetrical DEGs on SEQC data using each normalization method. The figure shows the ROC performance on RNA-Seq data with 733 PCR-validated genes.

**Fig. 3.**
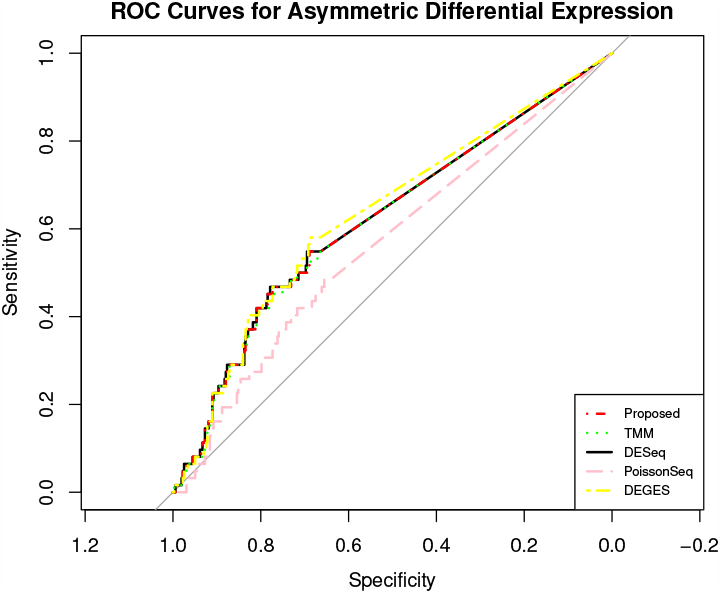
ROC curves for asymmetrical DEGs on SEQC data using each normalization method. The figure shows the ROC performance on RNA-Seq data with 619 PCR-validated genes.

The Microarray Quality Control Project (MAQC) data have two datasets, i.e., MAQC2, which has two replicates, and MAQC3, which has five replicates. The genes with low counts are filtered out similarly to the first datasets using the edgeR function filterByExpr. The filtered data is normalized using different normalization methods and DEs, and statistical tests are performed. The (Fig. 4 and 6) show the AUC curve of different approaches. The average AUC values are also presented (Table 2), which shows that the proposed method is better than the rest. We also examined the performance of the presented method on the two-group Pickrell dataset. After removing the genes with low counts, the data is normalized using all the methods. The DE results on the normalized data are 161 DE genes in the proposed approach, 162 in TMM, 53 in DESeq2, 48 in DEGES, and 45 in PoissonSeq. As shown (Fig. 5) on the Pickrell data, the AUC plot also indicates that the present approach can find better genes classified between the two classes than the other methods. The AUC values are also given (Table 2) for better comparison. We have also plotted the Venn diagram plot of the differentially expressed genes on the Pickrell dataset as shown (Fig. 7). The MA plot on the Pickrell dataset shows how many DE genes are identified by the proposed and state-of-the-art methods. The MA plot (Fig. 8) also shows the common DE genes between the present and other state-of-the-art methods.

**Table 2.** Comparison of AUC Values on Real Datasets with State-of-the-arts.

| Datasets |  | DESeq<br>(Love et al. [2014]) | DEGES<br>(Kadota et al. [2012]) | PoissonSeq<br>(Li et al. [2012]) | TMM<br>(Robinson and Oshlack [2010]) | Proposed |
| --- | --- | --- | --- | --- | --- | --- |
| Real Data | SEQC (SDE) | 79.38 | 89.08 | 66.02 | 83.69 | **91.52** |
|  | SEQC (ADE) | 62.09 | 63.25 | 56.67 | 61.81 | **62.01** |
|  | MAQC2 | 86.81 | 86.79 | 86.68 | 91.30 | **91.31** |
|  | MAQC3 | 91.47 | 91.10 | 91.16 | 94.07 | **94.98** |
|  | Pickrell | 91.66 | 92.01 | 92.34 | 91.56 | **93.66** |

SDE: Symmetric differential expression, ADE: Asymmetric differential expression

**Fig. 4.**
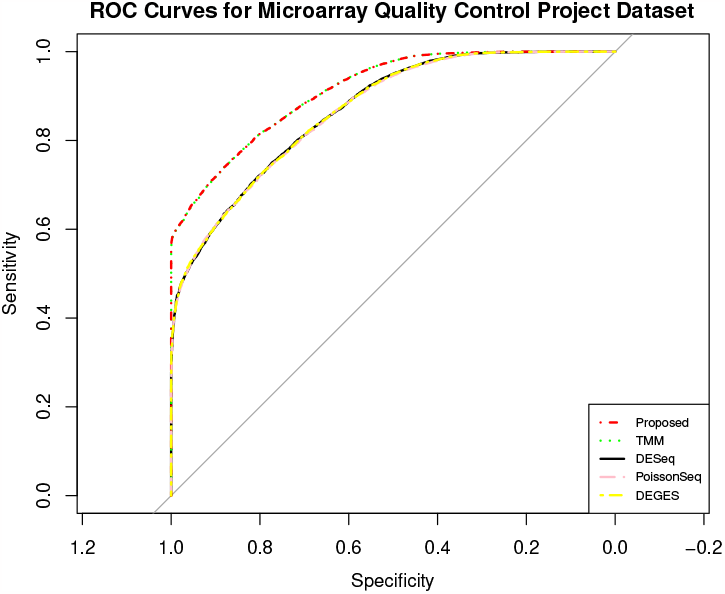
The ROC curves showing the performance of five normalization methods on MAQC2 dataset with two technical replicates in two-group condition.

**Fig. 5.**
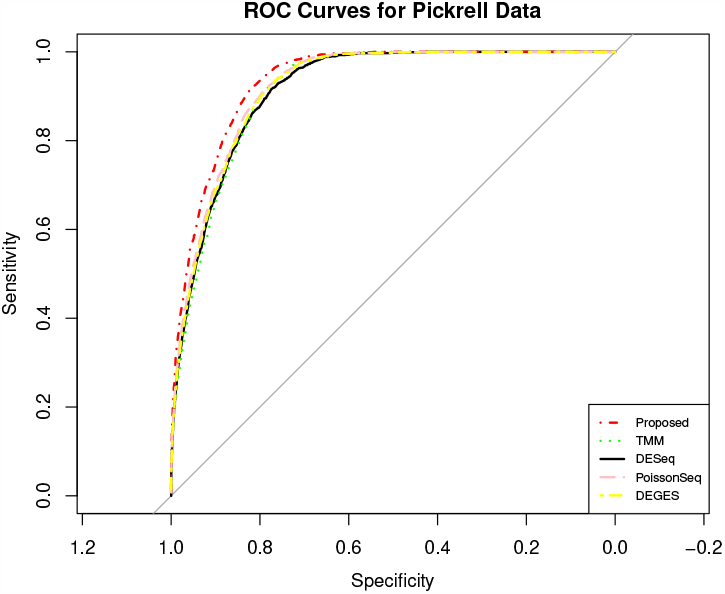
The ROC curves describing the performance of five normalization methods on two-group condition of Pickrell data.

**Fig. 6.**
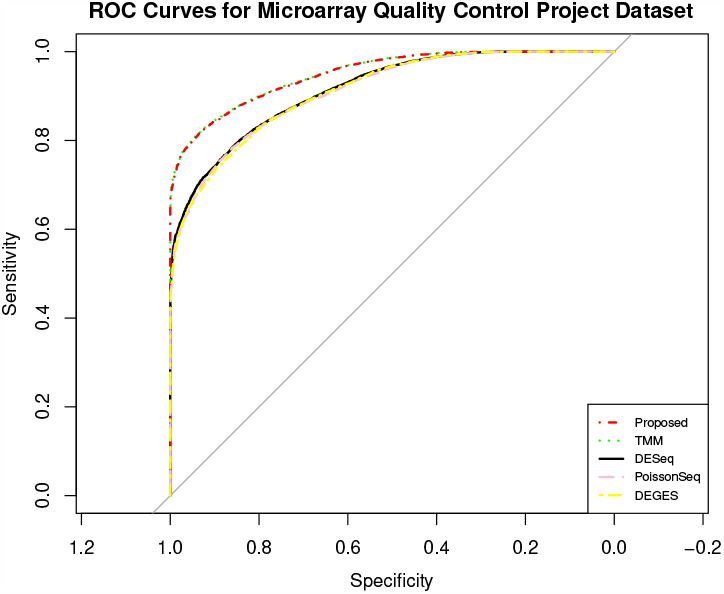
The ROC curves showing the performance of five normalization methods on MAQC3 dataset with five technical replicates in two-group condition.

**Fig. 7.**
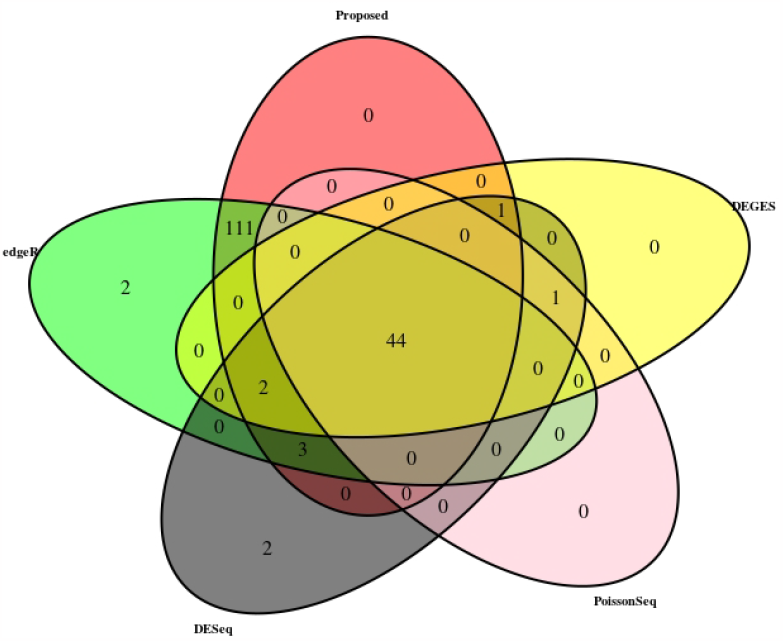
Venn Diagram on Pickrell dataset with pval*<* 0.01 & abs(log-fold-change)*>* 1.

**Fig. 8.**
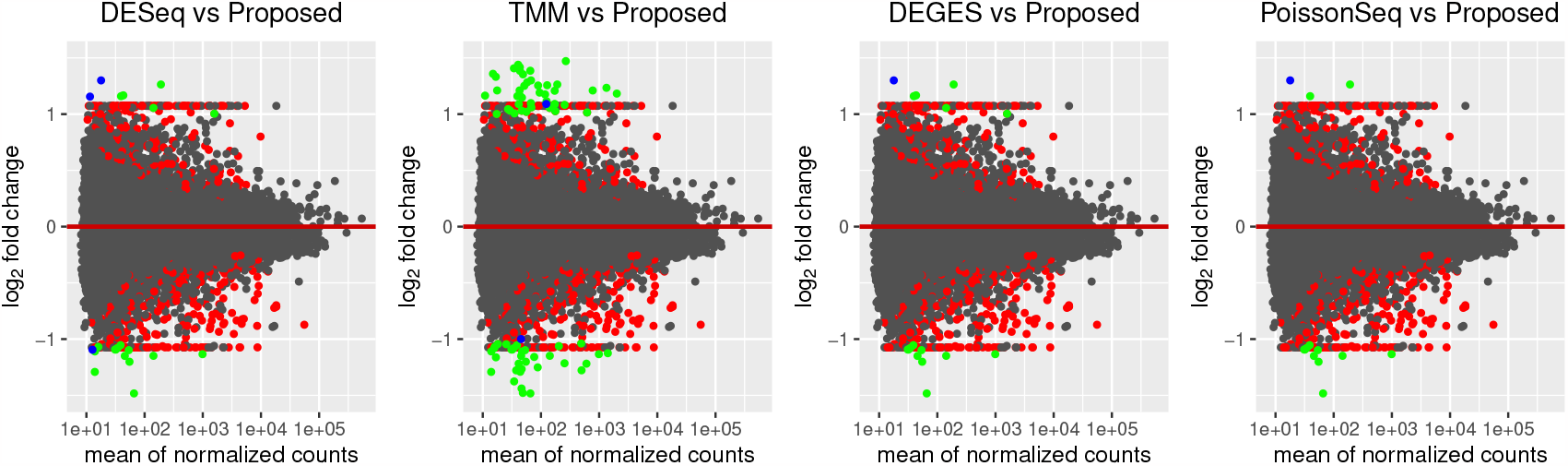
The red dots indicate DE genes identified only by proposed method. Then green dots are the shared results of proposed method and other state-of-the-art methods. The blue dots are DE genes identified only by state-of-the-art on the Pickrell dataset.

#### Performance Analysis on Simulated Datasets

The performance of the proposed approach has been experimented on the simulated datasets with two- and three-group conditions in terms of AUC values by varying the percentage of DEGs and up-regulated genes using the TCC function simulatedReadCounts. The AUC value provides data comparisons without a trade−off in specificity and sensitivity. The ROC curve depicts the true positive rate (i.e., sensitivity) vs the false positive rate (1− specificity) obtained for each threshold condition. In the two-group comparison we have varied the DEGs from 5%, 15% and 35%, repsectviely and up-regulated genes are varied from 5%, 15%, 25%and 35% as shown (Fig. 9, 10 and 11). However, in the case of three-group generated data, we have varied the DEGs from 5%, 15%, 25%, 35%, 55%, and 65% respectively, with proportion of up-regulated genes varied from 33.33% in each group as shown (Fig. 12) and 40%, 20% and 40% as shown (Fig. 13). The present approach performs better due to the variable trimming percentage of log-fold change.

**Fig. 9.**
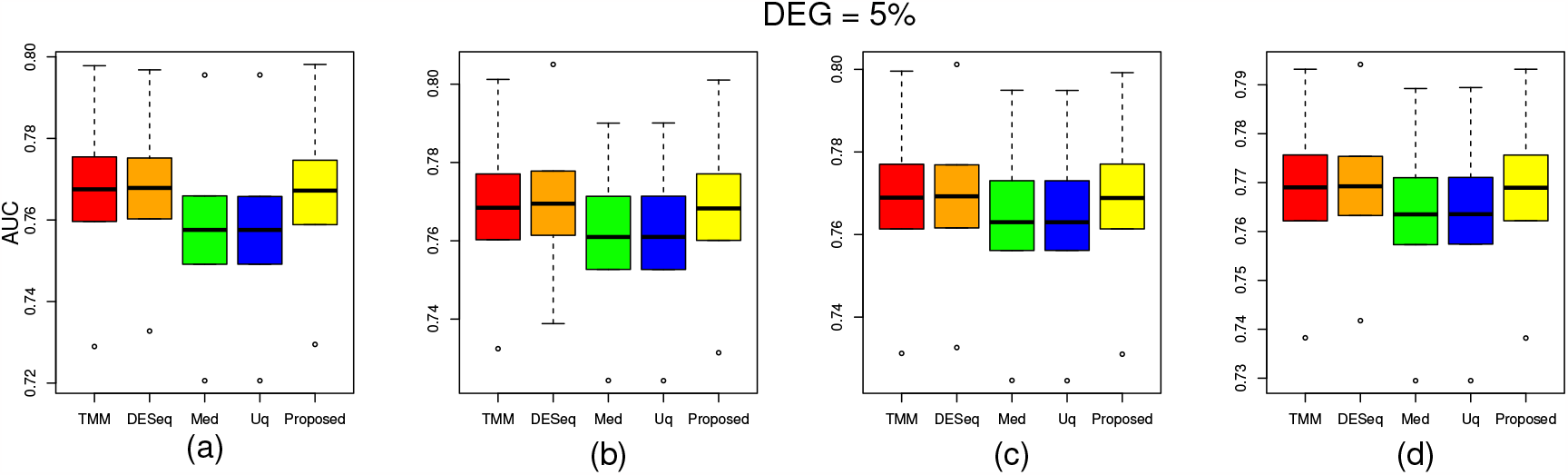
Two group simulated data have 5% deferentially expressed genes and each groups have three replicate with up-regulated deferentially expressed genes are varied from (a) 5%, (b) 15%, (c) 25%, (d) 35% in each groups

**Fig. 10.**
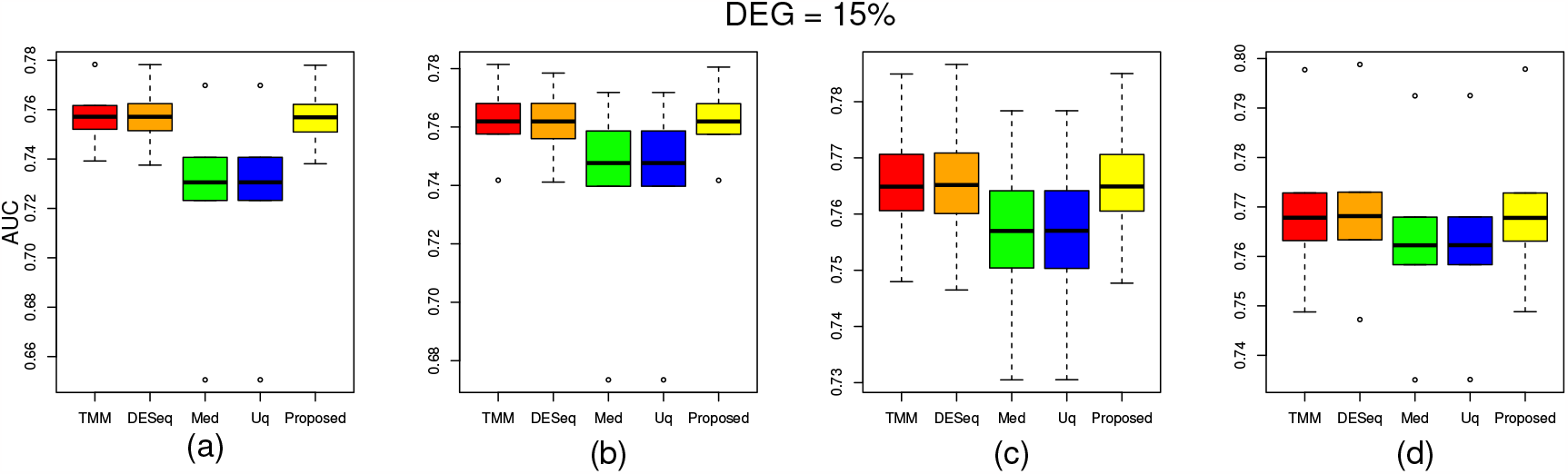
Two group simulated data have 15% deferentially expressed genes and each groups have three replicate with up-regulated deferentially expressed genes are varied from 5%, (b) 15%, (c) 25%, (d) 35% in each groups

**Fig. 11.**
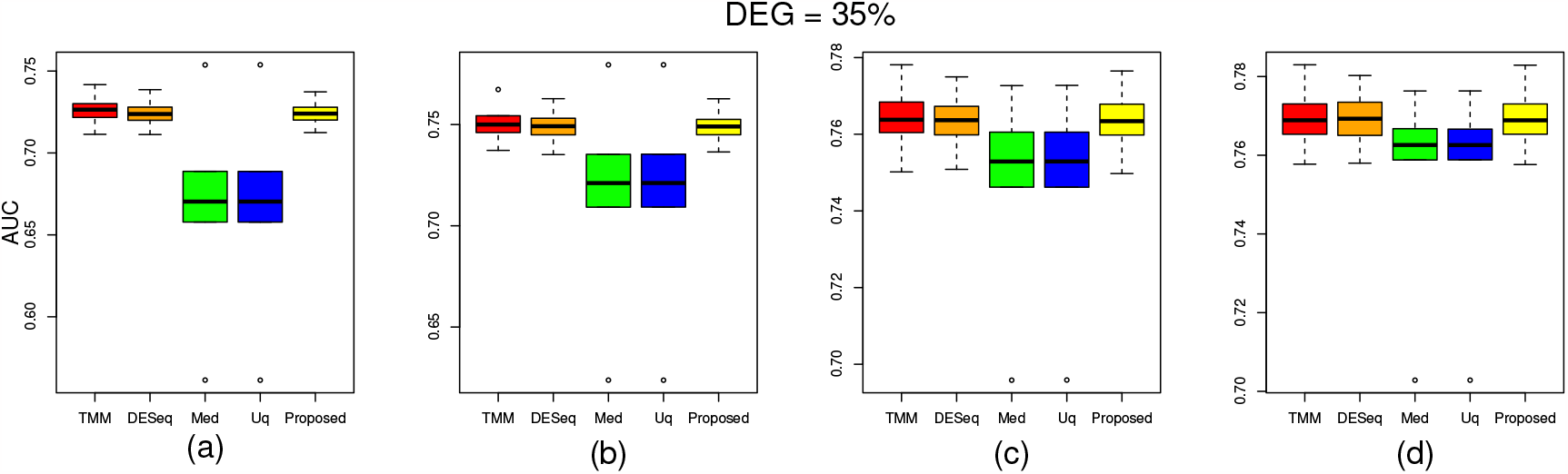
Two group simulated data have 35% deferentially expressed genes and each groups have three replicate with up-regulated deferentially expressed genes are varied from (a) 5%, (b) 15%, (c) 25%, (d) 35% in each groups

**Fig. 12.**
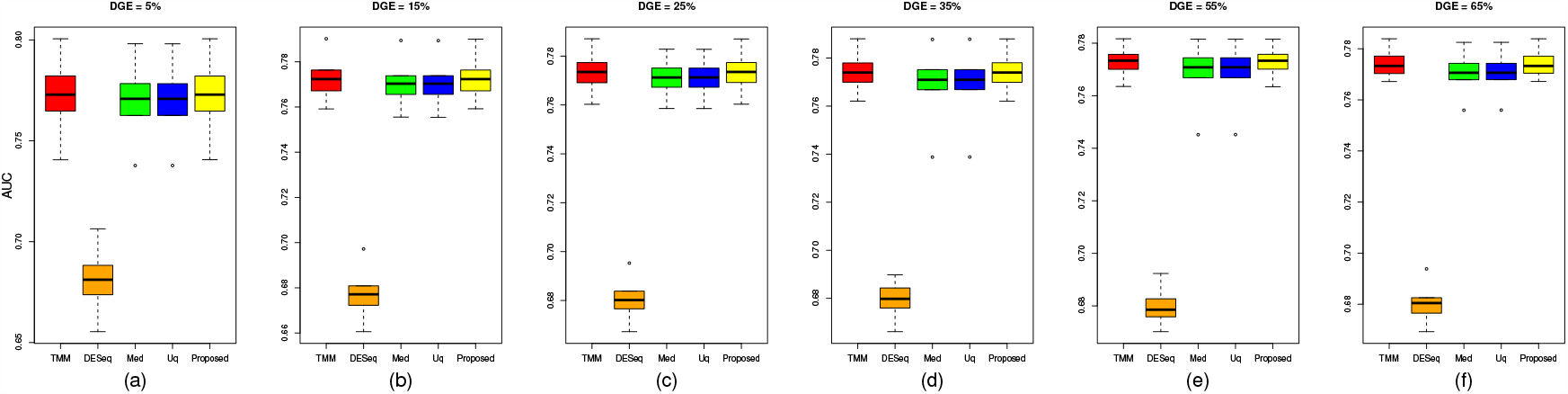
Three group simulated data with deferentially expressed genes are (a) 5%, (b) 15%, (c) 25%, (d) 35%, (e) 55%, (f) 65% and each groups have three replicate with up-regulated deferentially expressed genes are 33.33% in each groups respectively..

**Fig. 13.**
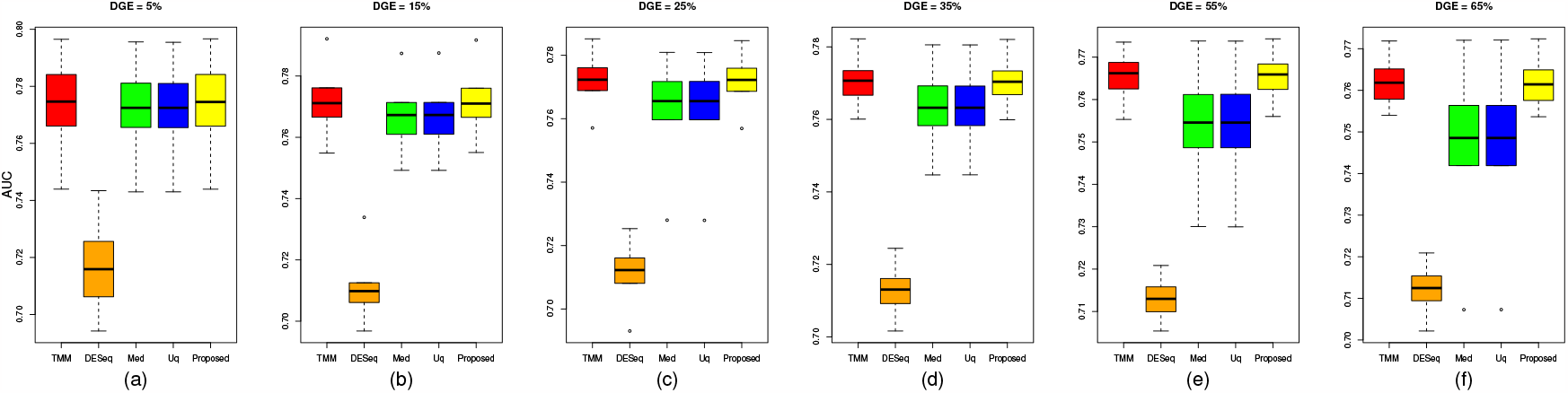
Three group simulated data with deferentially expressed genes are (a) 5%, (b) 15%, (c) 25%, (d) 35%, (e) 55%, (f) 65% and each groups have three replicate with up-regulated deferentially expressed genes are 40% in group 1, 20 % in group 2 and 40% in group 3 respectively.

**Fig. 14.**
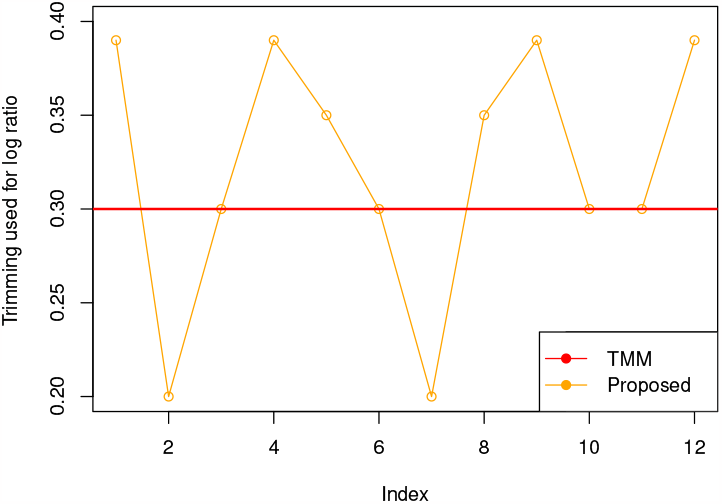
Trimming factor values are plotted for the MAQC2 dataset with proposed approach (orange line) with TMM (red line).

#### Trimming Factor

The trimming factor is an essential parameter to trim the data while preserving the desired information for the DEGs analysis. It is estimated using Jaeckel’s Estimator and plotted (Fig.14). As shown in the figure, we are getting different trimming values (change from 0.20 to 0.39 in case of MAQC2 dataset) for log fold change compared to TMM (by default, 0.30), which helps to find a better scaling factor while normalizing the samples.

## Conclusion

This paper presents an adaptive approach for bulk RNA-Seq data normalization. It automatically selects the trimming value of the *M* corresponding to the log fold change and absolute mean expression of RNA-Seq data to calculate the size factor for normalizing the data. It validated the real and simulated datasets to identify their effectiveness compared to state-of-the-art methods. The table 2 shows the effectiveness of the present approach on the real datasets. In the simulated datasets, as we varied the percentage of DE genes and up-regulated DE genes in the two and three-group conditions, the proposed method performed better, as described in the performance evaluation subsection and (Fig. 9, 10, 11, 12, and 13), respectively. The effectiveness in terms of performance shows that the present approach scales the datasets better than the rest of the methods in most cases since it trims each sample with different trimming values. The proposed approach will be extended to single-cell dataset normalization in the future.

## References

J Zyprych-Walczak, A Szabelska, L Handschuh, K Górczak, K Klamecka, M Figlerowicz, I Siatkowski, et al. The impact of normalization methods on RNA-Seq data analysis. BioMed research international, 2015, 2015.

Ciaran Evans, Johanna Hardin, and Daniel M Stoebel. Selecting betweensample RNA-Seq normalization methods from the perspective of their assumptions. Briefings in Bioinf., 19(5):776–792, 2018.

Ali Mortazavi, Brian A Williams, Kenneth McCue, Lorian Schaeffer, and Barbara Wold. Mapping and quantifying mammalian transcriptomes by RNA-Seq. Nature methods, 5(7):621–628, 2008.

Alicia Oshlack and Matthew J Wakefield. Transcript length bias in RNA-Seq data confounds systems biology. Biology direct, 4:1–10, 2009.

Davide Risso, Katja Schwartz, Gavin Sherlock, and Sandrine Dudoit. GC-content normalization for RNA-Seq data. BMC Bioinf., 12:1–17, 2011.

Vikas Singh, Nishchal K Verma, and Yan Cui. Type-2 fuzzy pca approach in extracting salient features for molecular cancer diagnostics and prognostics. IEEE Transactions on Nanobioscience, 18(3):482–489, 2019.

Vikas Singh and Nishchal K Verma. Gene expression data analysis using feature weighted robust Fuzzy-Means clustering. IEEE Trans. Nanobiosci., 22(1):99–105, 2022.

Taesung Park, Sung-Gon Yi, Sung-Hyun Kang, SeungYeoun Lee, Yong-Sung Lee, and Richard Simon. Evaluation of normalization methods for microarray data. BMC Bioinf., 4(1):1–13, 2003.

James H Bullard, Elizabeth Purdom, Kasper D Hansen, and Sandrine Dudoit. Evaluation of statistical methods for normalization and differential expression in mRNA-Seq experiments. BMC Bioinf., 11(1):1–13, 2010.

Cole Trapnell, Brian A Williams, Geo Pertea, Ali Mortazavi, Gordon Kwan, Marijke J Van Baren, Steven L Salzberg, Barbara J Wold, and Lior Pachter. Transcript assembly and quantification by RNA-Seq reveals unannotated transcripts and isoform switching during cell differentiation. Nature Biotechnol., 28(5):511–515, 2010.

Davide Risso. EDASeq: Exploratory data analysis and normalization for RNA-Seq. R package version, 1(0), 2011.

Mark D Robinson, Davis J McCarthy, and Gordon K Smyth. edgeR: a bioconductor package for differential expression analysis of digital gene expression data. bioinformatics, 26(1):139–140, 2010.

Michael I Love, Wolfgang Huber, and Simon Anders. Moderated estimation of fold change and dispersion for RNA-Seq data with DESeq2. Genome biology, 15(12):1–21, 2014.

Mark D Robinson and Alicia Oshlack. A scaling normalization method for differential expression analysis of RNA-Seq data. Genome biology, 11(3): 1–9, 2010.

Koji Kadota, Tomoaki Nishiyama, and Kentaro Shimizu. A normalization strategy for comparing tag count data. Algorithms for Molecular Biology, 7 (1):1–13, 2012.

Jun Li, Daniela M Witten, Iain M Johnstone, and Robert Tibshirani. Normalization, testing, and false discovery rate estimation for RNA-sequencing data. Biostatistics, 13(3):523–538, 2012.

Zhaonan Sun and Yu Zhu. Systematic comparison of rna-seq normalization methods using measurement error models. Bioinformatics, 28(20):2584–2591, 2012.

Vanessa M Kvam, Peng Liu, and Yaqing Si. A comparison of statistical methods for detecting differentially expressed genes from RNA-Seq data. American journal of botany, 99(2):248–256, 2012.

Charlotte Soneson and Mauro Delorenzi. A comparison of methods for differential expression analysis of RNA-Seq data. BMC Bioinf., 14(1):1–18, 2013.

Fatemeh Seyednasrollah, Asta Laiho, and Laura L Elo. Comparison of software packages for detecting differential expression in RNA-Seq studies. Briefings in Bioinf., 16(1):59–70, 2015.

Xiaohong Li, Guy N Brock, Eric C Rouchka, Nigel GF Cooper, Dongfeng Wu, Timothy E O’Toole, Ryan S Gill, Abdallah M Eteleeb, Liz O’Brien, and Shesh N Rai. A comparison of per sample global scaling and per gene normalization methods for differential expression analysis of RNA-Seq data. PloS one, 12(5):e0176185, 2017.

Stephen M Stigler. The asymptotic distribution of the trimmed mean. The Annals of Statistics, pages 472–477, 1973.

Remzi Oten and Rui JP de Figueiredo. Adaptive alpha-trimmed mean filters under deviations from assumed noise model. IEEE Trans. Image Processing, 13(5):627–639, 2004.

Z Su and CE Mason. SEQC/MAQC-III consortium a comprehensive assessment of RNA-Seq accuracy, reproducibility and information content by the sequencing quality control consortium. Nat. Biotechnol., 32(9): 903–914, 2014.

Leming Shi, Laura H Reid, Wendell D Jones, Richard Shippy, Janet A Warrington, Shawn C Baker, Patrick J Collins, Francoise De Longueville, Ernest S Kawasaki, Kathleen Y Lee, et al. The microarray quality control (MAQC) project shows inter-and intraplatform reproducibility of gene expression measurements. Nat. Biotechnol., 24(9):1151–1161, 2006.

Lin Wan and Fengzhu Sun. CEDER: accurate detection of differentially expressed genes by combining significance of exons using RNA-Seq. IEEE/ACM Trans. Comput. Biol. Bioinf., 9(5):1281–1292, 2012.

Franck Rapaport, Raya Khanin, Yupu Liang, Mono Pirun, Azra Krek, Paul Zumbo, Christopher E Mason, Nicholas D Socci, and Doron Betel. Comprehensive evaluation of differential gene expression analysis methods for RNA-Seq data. Genome biology, 14(9):1–13, 2013.

Leonardo Collado-Torres, Abhinav Nellore, Kai Kammers, Shannon E Ellis, Margaret A Taub, Kasper D Hansen, Andrew E Jaffe, Ben Langmead, and Jeffrey T Leek. Reproducible RNA-Seq analysis using recount2. Nat. Biotechnol., 35(4):319–321, 2017.

Joseph K Pickrell, John C Marioni, Athma A Pai, Jacob F Degner, Barbara E Engelhardt, Everlyne Nkadori, Jean-Baptiste Veyrieras, Matthew Stephens, Yoav Gilad, and Jonathan K Pritchard. Understanding mechanisms underlying human gene expression variation with RNA sequencing. Nature, 464(7289):768–772, 2010.

Sonja Hänzelmann, Robert Castelo, and Justin Guinney. GSVA: gene set variation analysis for microarray and RNA-Seq data. BMC Bioinf., 14:1–15, 2013.

Jianqiang Sun, Tomoaki Nishiyama, Kentaro Shimizu, and Koji Kadota. TCC: an R package for comparing tag count data with robust normalization strategies. BMC Bioinf., 14(1):1–14, 2013.

Min Tang, Jianqiang Sun, Kentaro Shimizu, and Koji Kadota. Evaluation of methods for differential expression analysis on multi-group RNA-Seq count data. BMC Bioinf., 16(1):1–14, 2015.

Takayuki Osabe, Kentaro Shimizu, and Koji Kadota. Differential expression analysis using a model-based gene clustering algorithm for RNA-Seq data. BMC Bioinf., 22(1):1–20, 2021.

